# Expansion Microscopy of Lipid Membranes

**DOI:** 10.1101/829903

**Authors:** Emmanouil D. Karagiannis, Jeong Seuk Kang, Tay Won Shin, Amauche Emenari, Shoh Asano, Leanne Lin, Emma K. Costa, IMAXT Grand Challenge Consortium, Adam H. Marblestone, Narayanan Kasthuri, Edward S. Boyden

## Abstract

Lipids are fundamental building blocks of cells and their organelles, yet nanoscale resolution imaging of lipids has been largely limited to electron microscopy techniques. We introduce and validate a chemical tag that enables lipid membranes to be imaged optically at nanoscale resolution via a lipid-optimized form of expansion microscopy, which we call membrane expansion microscopy (mExM). mExM, via a novel post-expansion antibody labeling protocol, enables protein-lipid relationships to be imaged in organelles such as mitochondria, the endoplasmic reticulum, the nuclear membrane, and the Golgi apparatus. mExM may be of use in a variety of biological contexts, including the study of cell-cell interactions, intracellular transport, and neural connectomics.

## Main

Expansion microscopy (ExM) physically magnifies biological specimens by covalently anchoring biomolecules or labels to a swellable polymer network (typically sodium polyacrylate) synthesized in situ throughout the specimen^1–4^. Following tissue softening and solvent exchange, the hydrogel-specimen composite expands isotropically, typically to a physical magnification of ~4.5x in linear dimension. The net result is that biomolecules or labels that are initially localized within the diffraction limit of a traditional optical microscope can now be separated in space to distances far enough that they can be resolved on ordinary microscopes. Expansion microscopy protocols^5^ for the visualization of proteins^3,6,7^ and nucleic acids^4^ are in increasingly widespread use, raising the question of whether other biological molecule classes, such as lipids, can also be visualized by ExM. We here report an expansion microscopy-compatible lipid stain, as well as a form of expansion microscopy (membrane expansion microscopy, or mExM) optimized for the imaging of lipid membranes as well as the antibody labeling of protein targets.

Traditional fluorescent membrane labeling probes (i.e., DiI and DiO) consist of long hydrophobic chains bearing fluorophores^8^, which preferentially localize to and diffuse within membranes^9^. Recent iterations of such molecules (i.e., FM1-43FX^10^ and mCLING^11^) also include hydrophilic moieties, like primary amines, to permit them to diffuse more freely in three dimensions throughout tissues prior to reaching their membrane targets. Inspired by these examples, which are not directly compatible with expansion microscopy, we sought to develop a membrane probe that is compatible with ExM chemistry, and in particular one that can achieve dense enough membrane coverage to support nanoscale resolution imaging and allow continuous tracing of membraneous structures (which degrades when stain density is insufficiently high). To achieve this, we designed a stain with the following features: (1) amphiphilicity, to support both lipid membrane intercalation and diffusion in 3D tissues, (2) a chemical handle for chemoselective conjugation of a fluorophore following the formation of the expandable hydrogel network, so that the molecular weight of the initial label can be kept as small as possible to facilitate diffusion, and (3) a polymer-anchorable handle for binding the probe to the ExM-gel matrix, to allow expansion.

We designed our membrane probe (Fig. 1a, lower left) to contain a chain of lysines, since they contain primary amines on their side chains that could serve as sites for binding to a polymer-anchorable handle, such as acryloyl–X (AcX), previously used to attach proteins via their amines to the in situ synthesized ExM hydrogel^3^. The amines are also positively charged, which can help promote interactions with negatively charged membranes^12^. To achieve lipid membrane intercalation, we included a lipid tail on the amine terminus of the lysine chain, with a glycine in between to provide mechanical flexibility^13^. Such glycine linkers are encountered next to lipid tails in many post-translationally modified proteins^14^. We limited the size of the label to ~1kDa, to allow fast diffusion throughout tissue and dense labeling of membranes; this limited us to having ~5 lysines in the backbone. We chose to use D-lysines, rather than the biologically typical L-lysines, to minimize degradation during the mechanical softening step of ExM, which in its most popular form involves a strong proteinase K digestion step. Finally, we introduced a chemical handle to the carboxy terminus of the terminal lysine. We chose to use a biotin, which could then react with one of the four binding sites of a fluorescently labeled streptavidin administered later^15^; the remaining active sites of the streptavidin can then be reacted with further biotinylated fluorophores to amplify the brightness manyfold.

**Figure 1.**
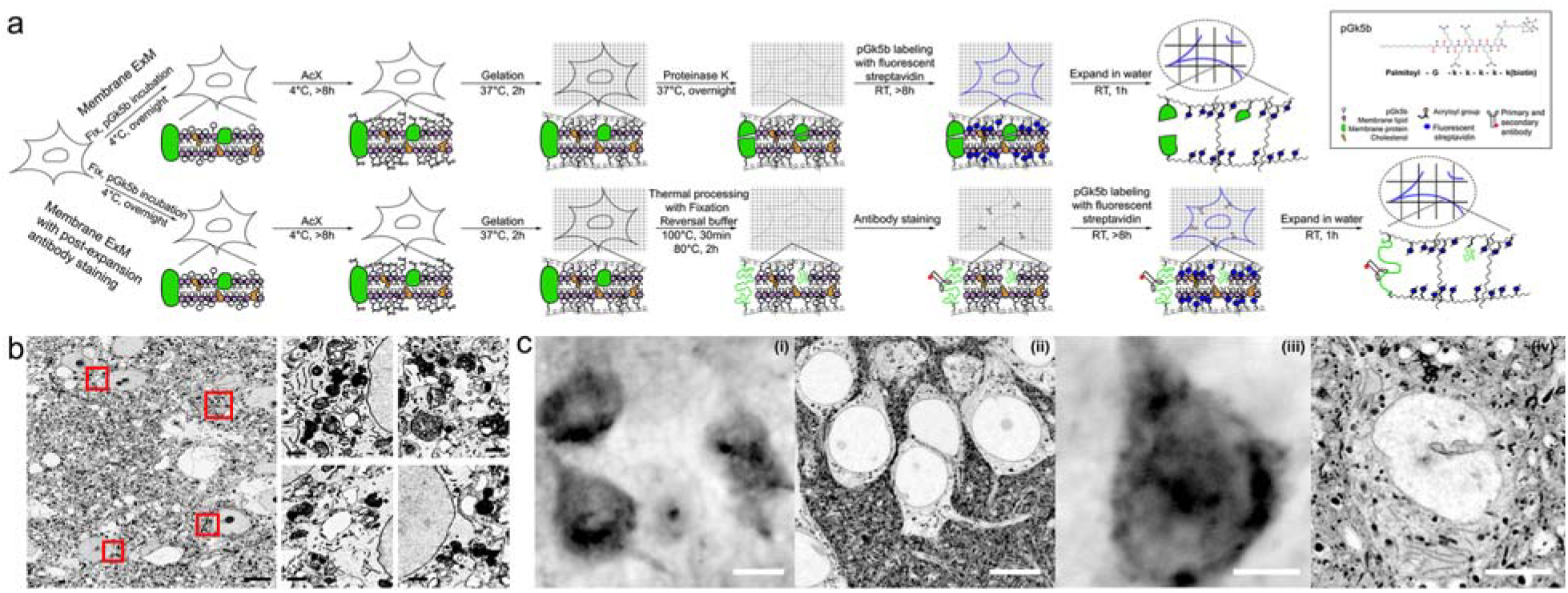
Membrane expansion microscopy (mExM) workflow and validation. (a) Tissue perfused and fixed with cold 4% PFA and 0.1% glutaraldehyde is sliced. Each tissue slice is incubated with 100μM of pGk5b membrane label (chemical structure, lower left) in PBS at 4□C overnight. The tissue is gelled following the established proExM protocol. If only membrane labeling is required, downstream processing of the tissue is similar to the already published proExM method^3^: the tissue is homogenized with proteinase K and labeled with fluorescent streptavidin (top row). pGk5b is not digested during proteinase K treatment because it is composed of D-amino acids. To incorporate antibody labeling, we developed a post-expansion antibody labeling technique (bottom row). In that case, the gelled tissue is thermally processed and expanded in a fixation reversal buffer, washed, and labeled with antibodies prior to the fluorescent labeling of pGk5b with streptavidin. (b) We validated the labeling efficacy of pGk5 to label membranes with electron microscopy. 100 μM of pGk5 (with azide instead of biotin) was applied to 200μm thick tissue slices from a mouse brain perfused with 4% PFA and 0.1% glutaraldehyde at 4□C, labelled with lipids for 2 days at 4□C, and labelled with 0.8nm undecagold gold nanoparticles conjugated to dibenzocyclooctyne. The tissue was post-fixed in 2% glutaraldehyde, embedded in resin, counter-labeled with osmium tetraoxide and imaged. When labeling membranes with the palmitoylated version of the probe (pGk5), the staining is dominated by the gold nanoparticles which overlap with osmium. The osmium staining (less contrast than the nanoparticles) becomes apparent when the tissue is labeled with the farnesylated probe, which exhibits less membrane coverage than the palmitoylated one (**Supp. Fig. 1**). Scale bars: 10 μm for the low-resolution images, 1 μm for the high-resolution insets. (c) Tissue labeled with pGk5b and unexpanded (i, zoomed out; iii, zoomed in) and post-expansion (ii, zoomed out; iv, zoomed in). All images were acquired with an Andor spinning-disk (CSU-W1 Tokogawa) confocal system on a Nikon Eclipse Ti-E inverted microscope body with a 40x 1.15 NA water-immersion objective. Either when observing a collection of neurons (i and ii) or a single neuron (iii and iv), mExM yields more detail than can be observed in the unexpanded sample, even using the novel lipid stain. The unexpanded images are sized to match the biological scale of the expanded sample. Scale bars represented in pre-expansion units: 10μm for (i) and (ii), and 5μm for (iii) and (iv).

We modified the preprocessing steps of the ExM protocol to better preserve lipids, since most lipids are not fixed during conventional chemical fixation^16^. To achieve lipid retention, we introduced two practices during tissue preservation and preparation for ExM. First, we combined paraformaldehyde with a low percentage (0.1%; increasing this number to 0.2% lowered the expansion factor, indicating incomplete tissue homogenization) of glutaraldehyde during fixation, since the latter helps to stabilize lipids (likely via stabilizing surrounding proteins^16^). Second, we maintained the temperature during tissue processing at 4□C (processing at room or higher temperatures decreased lipid signals), since lipid loss can be exacerbated by higher temperatures^17^, but at low temperatures lipids are in a more ordered state and are less likely to diffuse out of the sample^18^. In more detail: we preserved brain tissue in 4% paraformaldehyde (PFA) with 0.1% glutaraldehyde in ice cold phosphate buffered solution (PBS), sectioned the brain and washed out excess aldehydes, then applied the membrane probe (still at 4□C) at 100μM final concentration. Incubation at higher than 100μM probe concentrations yielded no change in the observed signal. We then added AcX at 4□C to make the membrane probes hydrogel-anchorable, incubated the specimen with the ExM gel monomer solution at 4□C, and finally incubated the samples at 37□C to initiate free-radical polymerization based hydrogel formation (**Supp. Table 1**). Finally, gelled samples were homogenized with a digestion buffer containing proteinase K, and post-labeled with fluorescent streptavidin (Fig. 1a, top row). Except for the pre-processing, the remainder of the steps – incubation, gelation, digestion, and expansion – are identical to earlier ExM protocols such as proExM, ensuring even and isotropic expansion.

We investigated, inspired by common lipid post-translational protein modifications, lipid tails that were both saturated and unsaturated -- specifically palmitoyl^19^ versus farnesyl^20^ tails. To assess whether saturated vs. unsaturated lipid tails exhibited different performance, we incubated mouse brain tissue sections with 100 μM of palmitoylated vs. farnesylated forms of our lipid stain (with an azide replacing the biotin, for simplicity). We post-labeled the lipid stains with 0.8 nm gold nanoparticles modified with a dibenzocyclooctyne (DBCO) handle, and imaged the resulting specimens with an electron microscope. We counterstained membranes with a “ground truth” electron microscopy label, osmium tetraoxide. These experiments showed that our construct indeed labeled lipids (Fig. 1b), and also that the palmitoylated probe (Fig. 1b) achieved denser membrane labeling than the farnesylated one (**Supp. Fig. 1**). This pattern was borne out when the stains were compared using ExM (**Supp. Fig. 2**). Thus, we chose palmitoyl as the lipid tail of our label. We tried combining a palmitoyl and a farnesyl group into a single backbone (on the N and C termini of the peptide, respectively), but observed limited tissue penetration, perhaps due to high hydrophobicity, as evidenced by probe accumulation on the surface of the tissue. We also assessed versions of the probe with vs. without the glycine linker; omitting the glycine linker caused loss of detail (**Supp. Fig. 3**). The results of these chemical synthesis and screening steps was a glycine and penta-lysine D-peptidic backbone, with a palmitoyl lipid group on the amine-terminus and a biotin on the carboxy-terminus, with a molecular weight of 1216 Daltons, which we call pGk5b (palmitoyl-G-kkkkk-biotin; Fig. 1a, inset). We call the use of this lipid label, with lipid-preservation-optimized tissue fixation and processing as described in the previous paragraph, membrane expansion microscopy (mExM). We have synthesized a significant amount of pGk5b and will distribute it, free of charge, to people interested in trying it out; we are also beginning the process, in parallel, of seeking a commercial vendor.

mExM enabled dense labeling of membranous structures in the mouse brain that cannot be observed in unexpanded tissue (**Fig.1c**). In addition to tissue specimens (Fig. 2), we can use mExM to label and expand membranes of fixed cells cultured in vitro (**Supp. Fig. 4**). Samples expand 4.5 times, as expected because the core (polymerization, digestion, and expansion) of the mExM process is identical to that of earlier ExM protocols such as proExM, which have been validated to expand isotropically (with few-percent error over length scales of tens to hundreds of microns) in many cell types and tissue types^3,21,22^.

**Figure 2.**
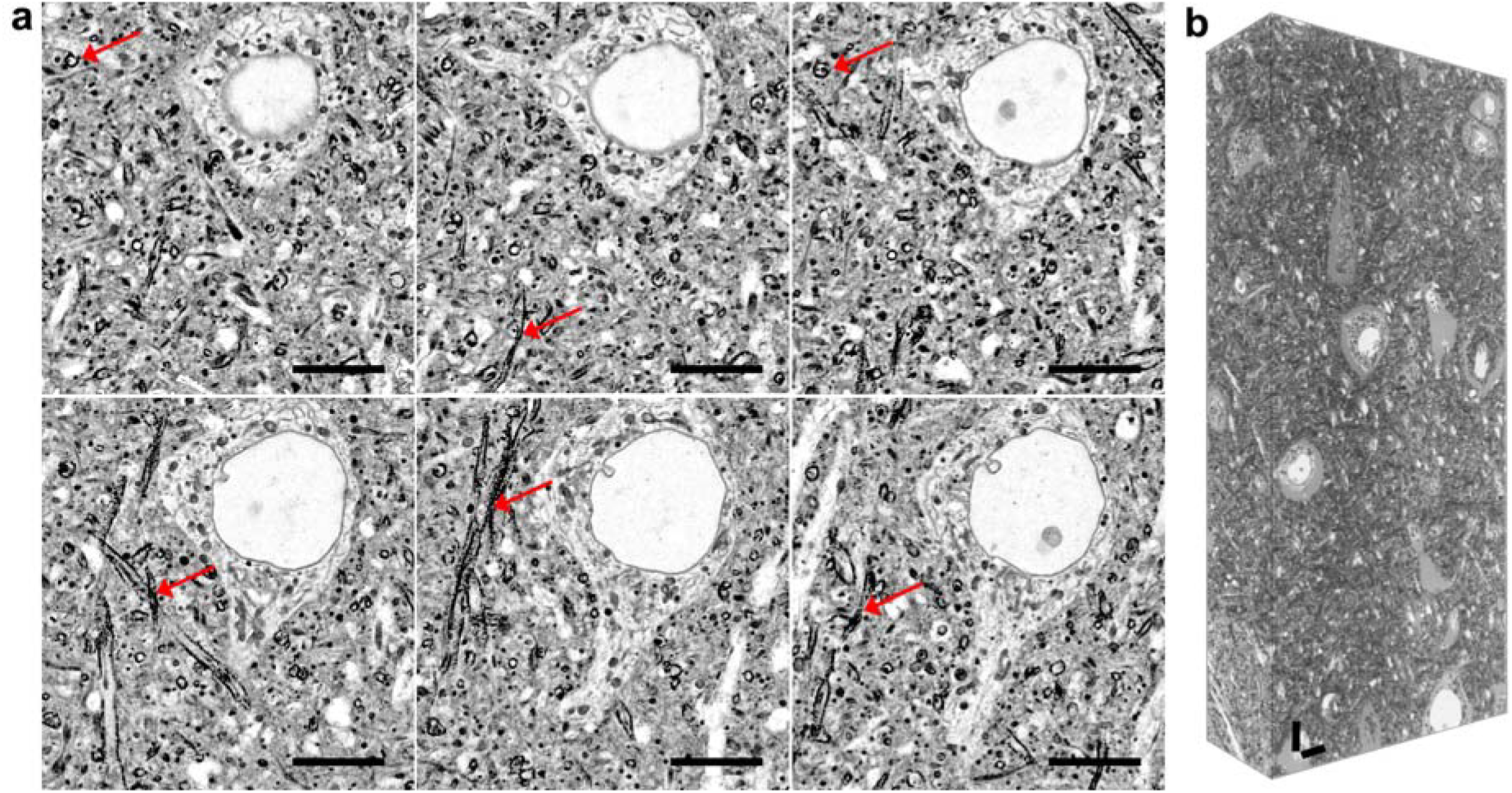
Membrane expansion microscopy (mExM) of fixed brain tissue. (a) mExM enables the labeling of membranes in thick pieces of mouse brain tissue. Here shown are six serial sections from a 3D image stack taken with a confocal spinning disk microscope. Axons can be identified by their high contrast due to the increased concentration of lipids in myelin (details shows with red arrows). (b) mExM processed tissue imaged with light-sheet microscopy. Scale bars represented in pre-expansion units: 10μm.

### Immunohistochemistry-compatible mExM

Given the interactions between proteins and lipids, in support of the scaffolding of cellular signals and compartmentalization of signaling cascades, it would be desirable to jointly image lipids and proteins in the same specimen. We therefore further developed an integrated protein and lipid ExM imaging protocol. Combining antibody staining and lipid labeling was not trivial: immunostaining protocols typically require permeabilization of samples with detergents, to enable penetration of antibodies throughout tissue^23^, but detergents will also solubilize lipids and permeabilize membranes, preventing retention of the nanoscale spatial layout of the lipid membranes^24^. We tried a variety of methods to resolve this issue. Using mild detergents (compared to those typical in immunofluorescence, e.g., Triton-X, NP-40, TWEEN) like saponin^25^ resulted in antibody signals that looked punctate and dim, and compromise details of intracellular organelles. Instead, we therefore sought to immobilize the lipid probes before permeabilization and antibody labeling. To achieve this, we initially attempted re-fixing the lipid probes by exposure to 0.1% PFA, but observed loss of epitope accessibility upon immunostaining, as well as low expansion factor (x2.8) (**Supp. Fig. 5a**). Alternatively, prior to forming the ExM gel, we formed a cleavable but non-expanding hydrogel (containing only uncharged acrylamide, and crosslinked with the cleavable N,N’-diallyl-tartardiamide), that could anchor the lipid labels in place prior to detergent exposure, antibody staining, gelation with an expanding gel, cleavage of the initial gel and finally expansion of the second gel. In this case, the lipid staining was preserved, but antibody labeling was dim -- presumably due to diffusion limitations arising from the non-expanding gel (**Supp. Fig. 5b**).

To solve these problems, we developed a *post-expansion* antibody labeling protocol, in which we first labeled the tissue lipids with pGk5b, synthesized the ExM gel, used high temperature and detergent (rather than epitope-destroying proteinase K) to perform the mechanical homogenization process^23^ which allows isotropic expansion^3^, and then delivered antibody labels to the retained epitopes after expansion. After this homogenization process but before replacing the buffer with pure water, to achieve full 4.5x expansion, the sample expands ~2 times in the antibody labeling buffer; thus we denote this procedure as “post-expansion antibody labeling”. Our new protocol builds on the post-expansion protein-retention ExM (proExM) protocol^3^, as well as tissue proteomics protocols for formalin-fixed paraffin-embedded (FFPE) tissues^26–28^. In brief, we heat the sample for half an hour at 100□C and for 2 hours at 80□C, in a “fixation reversal” (FR) buffer^28^ containing 0.5% PEG20000, 100mM DTT, 4% SDS, in 100mM Tris pH8 (**Supp. Table 2**). We found the FR buffer to work better than high-temperature treatment with alternative buffers (Citrate or Tris buffer for antigen retrieval) that caused loss of antibody staining (**Supp. Fig. 6**), or commercially available protein extraction buffers (i.e., RIPA buffer^29^ and the Liquid Tissue MS Protein Prep Kit^30^) that did not support tissue expansion. Alternatively, the samples can be autoclaved at 121□C for 1 hour in the FR buffer. In that case, and following the observations established for the post-expansion proExM method^3^, some epitopes were not stained effectively by antibodies (**Supp. Table 2**), however, perhaps because of an interaction between the FR buffer and very high temperatures.

We performed mExM with antibody staining (Fig. 1a, bottom row), using antibodies against organelle-specific membrane-localized proteins -- such as calnexin for the endoplasmic reticulum (ER, Fig. 3a), giantin for the Golgi apparatus (Fig. 3b), Tom20 for mitochondria (Fig. 3c), and NUP98 for the nuclear membrane (Fig. 3d). We also labeled myelin using an antibody against myelin basic protein (Fig. 3e). In all cases, we observed clear co-localization of the lipid signals from the membranes constituting the organelles, with the antibody signals from the organelle membrane proteins.

**Figure 3.**
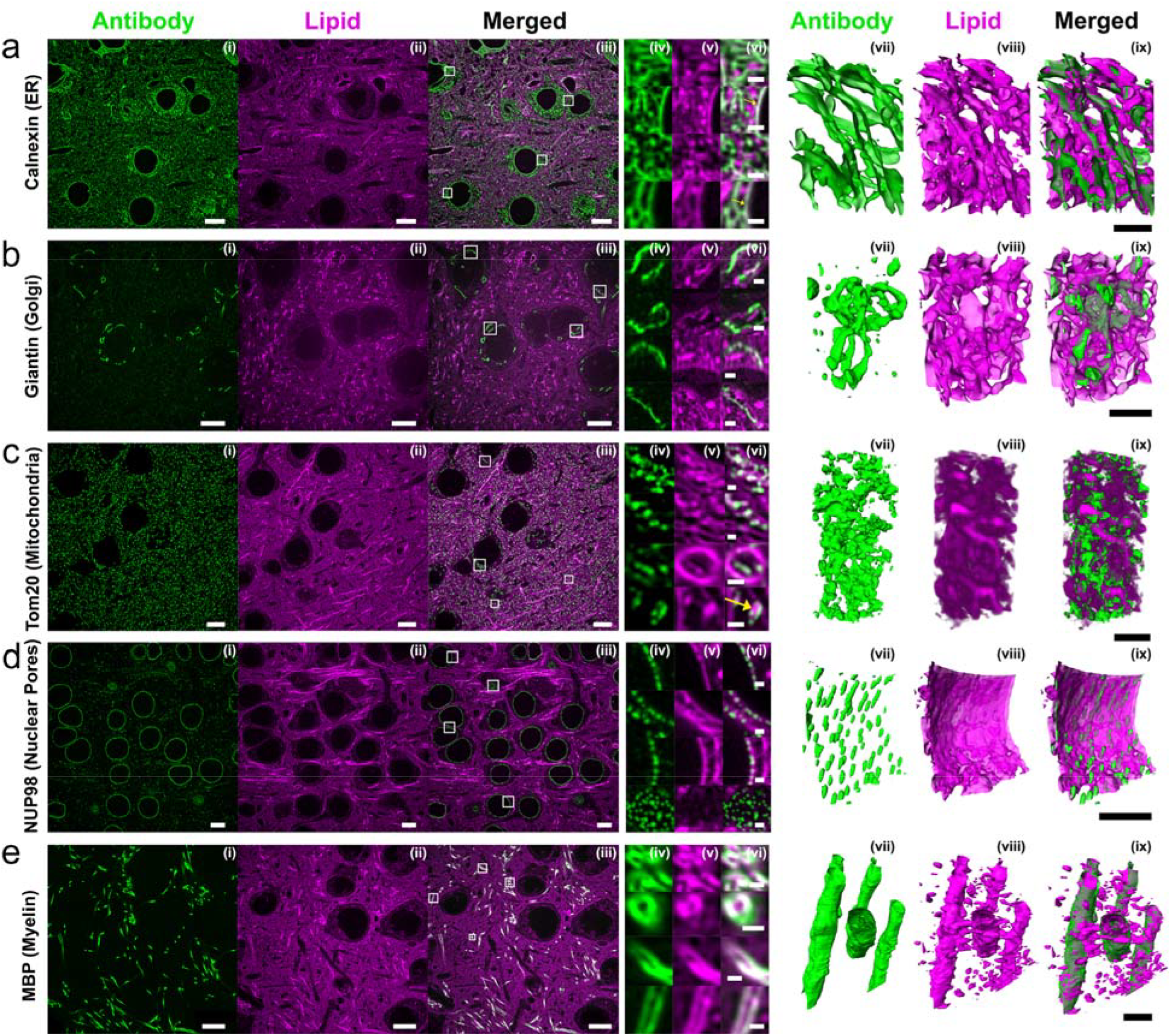
mExM enables simultaneous antibody and lipid co-visualization. After fixation and labeling with 100μΜ pGk5b, mouse brain tissue is gelled and processed in the FR buffer (0.5% PEG20000, 100mM DTT, 4% SDS, in 100mM Tris pH8) by heating at 100□C for 30min, then 80□C for 2hr. After washing in PBS the gels are labeled with antibodies for (a) an endoplasmic reticulum surface protein (Calnexin), (b) a Golgi apparatus marker (Giantin), (c) a mitochondrial membrane protein (Tom20), (d) a nuclear pore complex component (NUP98), and (e) myelin (Myelin Basic Protein). (i-ii) Individual signals for antibodies (green) and membranes (magenta) and (iii) the overlay of (i) and (ii). (iv-vi) Details from the regions of (iii) indicated by squares. (iv) Antibody signal (green), (v) membrane signal (magenta) and (vi) the overlay of (iv) and (v). (vii-ix) Spatial visualization in 3D (namely, stills from 3D movies contained in **Supp Movies 1-5**) of lipid and antibody co-labeling with mExM. The membrane label is in magenta and antibodies in green. (a) Calnexin stains for rough endoplasmic reticulum and co-localizes with the membrane signal. Calnexin also co-localizes with the nuclear membrane signal (yellow arrows in vi). (b) Giantin is expressed on the surface of the Golgi apparatus and also co-localizes with membrane signals. (c) Mitochondrial staining is prevalent throughout the tissue and the Tom20 antibody signal overlaps with membrane labels. Tom20 appears to cluster at the mitochondrial membrane (arrow in vi). (d) Nuclear pore complexes span throughout the nuclear membrane. (e) Myelin basic protein co-localizes with the membrane signal and exhibits dense labeling, corresponding to the amount of lipid in highly myelinated regions of axons. Scale bars represented in pre-expansion units: (i-iii) 10μm, (iv-vi) 1μm. (vii-ix) Scale bars represented in pre-expansion units: 5μm.

The high resolution of mExM captured details of protein-lipid organization known to occur, but difficult or impossible to assess with conventional microscopy. For example, it is known that the endoplasmic reticulum contains components that come in close apposition with the nuclear membrane^31^, which can be observed directly with ExM (Fig. 3a, right, yellow arrows). Specifically, Calnexin is a lectin protein expressed primarily on the surface of the rough ER^32^, and a 3D representation of the iso-intensity profiles of Calnexin protein expression in the tissue reveals the cisternae of the ER and its co-localization with mExM-tagged ER lipids^33^ (Fig. 3a(vii-ix) and **Supp. Movie 1**). Calnexin exhibits a peri-nuclear labeling pattern with co-localization with membrane labels – this represents a previously known sub-population of Calnexin proteins that are post-translationally modified with a palmitoyl group inside the ER and localize to contact sites of the ER with the nucleus^34^. Additionally, the Golgi apparatus, co-labeled with anti-giantin and our lipid stain, is visible primarily in neuronal somas, in accordance with previous studies^35^ (Fig. 3b, right). In 3D, giantin labeling exhibits cisternal morphology that overlaps with pGK5b labeling (Fig. 3b(vii-ix) and **Supp. Movie 2**). Mitochondria are abundant throughout the tissue in somata, dendrites and axons. (Fig. 3c(vii-ix) and **Supp. Movie 3**). Tom20 expression on the surface of mitochondria reveals individual protein clusters, consistent with other super-resolution imaging methods (Fig. 3c, right, Fig. 3c(vi) yellow arrow)^36^. Nuclear pore complexes are resolved as individual pores on the nuclear membrane (**Fig. 4d**, right), and can be seen to span the nuclear membrane when visualized in 3D (Fig. 3d(vii-ix) and **Supp. Movie 4**). Myelinated regions of neurons exhibit very strong membrane labeling (Fig. 3e, right), with excellent co-localization of the protein with the membrane label (Fig. 3e(vii-ix) and **Supp. Movie 5**).

In summary, we have developed a membrane intercalating probe that enables the imaging of cellular membranes, in thick fixed tissue, in the context of a lipid-optimized form of expansion microscopy. We also developed a post-expansion antibody labeling method that allows for the joint imaging of proteins and membranes in ExM. This enables nanoscale observation of lipid membrane structures and their associated proteins at nanoscale resolution, even across large volumes, using ordinary microscopes. We hope that this will democratize and extend the study of membrane conformation and signaling in a wide range of biological systems in normal and disease states. Future directions may include the development of an iterative^2^ form of mExM, in which a specimen is expanded twice (for a 4.5 × 4.5 ~20x physical magnification), which may enable resolutions of 10-20 nm, approaching that of electron microscopy, to be achieved.

## Acknowledgments

For funding, E.S.B. acknowledges Lisa Yang and Y. Eva Tan, John Doerr, the Open Philanthropy Project, the MIT Media Lab, the HHMI-Simons Faculty Scholars Program, the US Army Research Laboratory and the US Army Research Office under contract/grant number W911NF1510548, Cancer Research UK Grand Challenge grant C9545/A24042, the MIT Brain and Cognitive Sciences Department, the New York Stem Cell Foundation-Robertson Investigator Award, NIH Transformative Award 1R01GM104948, NIH Director’s Pioneer Award 1DP1NS087724, NIH 1R01EY023173, NIH 1U01MH106011, NIH 1R01EB024261, NIH 1R01NS102727, NIH 1R01MH110932, NIH 2R01DA029639, and the MIT McGovern Institute MINT program. J.S.K. acknowledges funding by the Samsung Scholarship. T.S. acknowledges funding by the National Science Foundation Graduate Research Fellowship Grant No. 1122374.

## Author Contributions

The study was devised by EDK, JSK and ESB with early input from AHM. Laboratory work was performed by EDK, JSK, TS, LW, EKC and NK. Image processing was performed by EDK and TS. Mouse perfusion and tissue preparation was performed by JSK, AE, LW and EKC. Spinning-disk microscopy imaging was performed by EDK, JSK and TS. Light-sheet microscopy imaging was performed by SA. ESB supervised the project. All authors contributed to the writing and editing of the manuscript.

## Materials and Methods

### Lipid Label Synthesis

The lipid labels were commercially synthesized (Anaspec). They were purified to >95% purity. They were aliquoted into 1 mg quantities in tubes, lyophilized to solid powder, and stored at – 20□C until stock solutions were prepared. For stock solutions, 1 mg of solid lipid label was dissolved in 50% DMSO and 50% ultrapure water to 10mM and stored in −20□C.

### Brain Tissue Preparation

Wild type (C57BL/6, Taconic) mice were first terminally anesthetized with isoflurane. Then, 1x phosphate buffered saline (PBS) was transcardially perfused until the blood cleared. For all mExM experiments, the mice were then transcardially perfused with the fixative 4% paraformaldehyde (PFA) and 0.1% glutaraldehyde, buffered in 1x PBS. The fixative was kept on ice during perfusion. After the perfusion step, brains were dissected out, stored in the fixative at 4□C overnight for further fixation, and sliced on a vibratome (Leica VT1000S) at 100 μm or 200 μm thickness. The slices were then kept in 1x PBS at 4□C overnight for washing and storing.

All procedures involving animals were in accordance with the US National Institutes of Health Guide for the Care and Use of Laboratory Animals and approved by the Massachusetts Institute of Technology Committee on Animal Care.

### mExM

Tissue slices (100 μm for light microscopy and 200 μm for electron microscopy) were first incubated in the lipid labels (e.g., pGk5b) at 4□C overnight to let the labels diffuse and intercalate thoroughly throughout the tissue slices. The lipid label 10mM stock solution was diluted in 1x PBS that was kept at 4□C, at 1:100 dilution, for incubating the tissue slices. For a single piece of tissue 1 ml of solution was prepared. Subsequently, 6-((acryloyl)amino)hexanoic Acid, Succinimidyl Ester (AcX) stock solution (10 mg/mL in dimethylsulfoxide (DMSO)) was diluted in 1x PBS at 1:100 dilution, and tissue slices were incubated overnight at 4□C. The AcX stock solution was prepared by dissolving 5 mg AcX (ThermoFisher, catalog no. A20770) in 500 μl anhydrous DMSO (ThermoFisher, catalog no. D12345). It is essential that both these steps are carried out at 4□C to keep the lipids in the tissue membranes thermodynamically stable. Then, the standard expansion microscopy steps were carried out. For formulations check **Supp. Table 1**. Briefly, for monomer solution we prepared Stock X and later added the polymerization initiator, accelerator and inhibitor. Recipe for Stock X: 8.6% (w/v) sodium acrylate (Sigma Aldrich, catalog no. 408220), 2.5% (w/v) acrylamide (Sigma Aldrich, catalog no. A8887-500G), 0.15% (w/v) N,N’-methylenebisacrylamide (Sigma Aldrich, catalog no. M7279-25G), 11.7% (w/v) sodium chloride (Thermo Fisher, catalog no. BP358-212), PBS). Recipes for the initiator, accelerator and inhibitor: 4-Hydroxy-TEMPO stock solution (4HT; Sigma Aldrich, catalog no. 176141), 0.5% (w/v) in water, N,N,N′,N′-Tetramethylethylenediamine stock solution (TEMED; Sigma Aldrich, catalog no. T7024-50ml), 10% (w/v) in water, and ammonium persulfate stock solution (APS; Thermo Fisher, catalog no. 17874), 10% (w/v) in water were prepared in advance and stored at −20C; for detailed step-by-step instructions, please visit: http://expansionmicroscopy.org/. The gelation solution was prepared by mixing Stock X, 4HT, TEMED, and APS stock solutions in a 47:1:1:1 ratio on a 4□C cold block, and the tissue slices that were incubated in the lipid labels and AcX were incubated in the gelation solution for 30 minutes at 4□C. During this step, the gelation chamber was constructed (**Supp. Fig. 7**). The chamber containing the tissue was then transferred over to an incubator kept at 37□C to initiate free-radical polymerization. After 2 hours, the gelation chamber containing the tissue was taken out, and the gelled tissue was cut out from the chamber to be immersed in proteinase K digestion buffer. Proteinase K (ProK; NEB, catalog no. P8107S) was stored at −20□C at 800U/ml concentration, which was then diluted in the digestion buffer at 1:100 concentration prior to use. The digestion buffer was prepared by mixing: Triton X-100 (Sigma Aldrich, catalog no. 93426) to a final concentration of 0.5% (w/v) in water, EDTA disodium (0.5 M, pH 8, Thermo Fisher, catalog no. 15575020) to final 1mM in water, Tris-HCl ((1 M) aqueous solution, pH 8, Life Technologies, catalog no. AM9855) to final 50mM in water, and Sodium Chloride (Thermo Fisher, catalog no. BP358-212) to final 1M in water. The gelled tissue was then digested at 37C on a shaker overnight. After digestion, the gelled sample was washed 4 times in 1x PBS at room temperature (RT), 30 minutes each. For imaging, the digested tissue was labelled with 0.3mg/ml of streptavidin labelled with Atto 565 (Atto 565-Streptavidin; Sigma Aldrich, catalog no. 56304-1MG-F) buffered in 1x PBS overnight at RT, and then washed 4 times 30 minutes each in 1x PBS at RT before it was placed in excess water for expansion. The fluorescence of lipid labels were further amplified by labelling the gel once more with biotin labelled with Atto 565 (Atto 565-Biotin; Sigma Aldrich, catalog no. 92637-1MG-F) at 0.01 mg/ml concentration after the gel was incubated in Atto 565-Streptavidin and before it was immersed in excess water for expansion. Alternatively, instead of using Atto 565-Streptavidin, wild type streptavidin (Thermo Fisher, catalog no. S888) was used at 0.3 mg/ml with the same labelling conditions, and Atto 565 biotin was added at 0.01mg/ml. The fluorescent labels attached to the biotin alone are sufficiently bright for imaging the sample with high signal-to-noise profiles.

### Immunohistochemistry-compatible mExM

The aforementioned mExM steps were carried out the same, except for the digestion step. Instead of using the proteinase K digestion buffer, tissue was boiled in the Fixation Reversal (FR) buffer for 30 minutes at 100□C and then held for 2 hours at 80□C, or autoclaved for 1 hour at 121□C. The FR buffer consists of 0.5% PEG20000, 100mM DTT, 4% SDS, in 100mM Tris pH8. After this, the FR-digested sample was washed in 1x PBS 4 times at RT for 1 hour before proceeding to the standard immunohistochemistry steps. The sample was first blocked with MAXblock Blocking Medium (Active Motif, catalog no. 15252) for 4-6 hours at room temperature and incubated in MAXbind Staining Medium (Active Motif, catalog no. 15251) containing primary antibodies at a concentration of 10 μg/mL overnight at 4□C. Then, the sample was washed with MAXwash Washing Medium (Active Motif, catalog no. 15254) at RT 4 times, 30 minutes each and subsequently incubated in secondary antibodies buffered in MAXbind Staining Medium at a concentration of 10 μg/mL for 10-12 hours at 4□C. Finally, the secondary antibodies were washed, again, with MAXwash Washing Medium at RT 4 times, 30 minutes each time. For primary antibodies, anti-Calnexin (Abcam, catalog no. ab22595, Rabbit), anti-Tom20 (Cell Signaling Technology, catalog no. 42406S, Rabbit), anti-NUP98 (Cell Signaling Technology, catalog no. 2597S, Rabbit), anti-Giantin (Biolegend, catalog no. 924302, Rabbit), anti-Myelin Basic Protein (MBP; Cell Signaling Technology, catalog no. 78896S, Rabbit), were used. For secondary antibodies, anti-Chicken Alexa Fluor Plus 488 (Thermo Fisher, catalog no. A32931), anti-Rabbit Alexa Fluor Plus 488 (Thermo Fisher, catalog no. A32731), and anti-Mouse Alexa Fluor Plus 647 (Thermo Fisher, A32728) were used. After antibody staining, the lipid labels that were conjugated to the gel were then labelled with streptavidin and biotin containing Atto 565 fluorophores, as described in the mExM protocol mentioned above.

### Confocal and Light-Sheet Imaging

All mExM images (Figure 2a, 3 and Supp. Figure 2, 3, 4, and 5) were taken with an Andor spinning disk (CSU-W1 Tokogawa) confocal system on a Nikon Eclipse Ti-E inverted microscope body with a 40x 1.15 NA water-immersion objective. Light-sheet imaging (Fig 2b) was performed with a Zeiss Z.1 Light-Sheet microscope, utilizing a 20x 1.0 NA water immersion lens.

### Expansion Factor Measurement

To determine the expansion factor for all mExM experiments, two procedures were followed. First, the distance between two landmarks in the whole tissue specimen was measured and compared before and after the expansion. Second, the tissue thickness, as measured by imaging the fluorescent labels across the vertical section of the specimen on a microscope, was determined and compared before and after expansion. The expansion factors that were obtained following these two ways were then averaged to be able to determine the expansion factor of an mExM experiment.

### Image Processing

Images shown in Figures 2-3 were first filtered by a Gaussian filter, and then processed by the CLAHE algorithm^37^ to remove possible noise. Image processing was performed with a 64GB RAM, 8 core CPU desktop.

