## Supplemental Material for "Expansion Microscopy of Lipid Membranes"

**Supplementary Figure 1.**

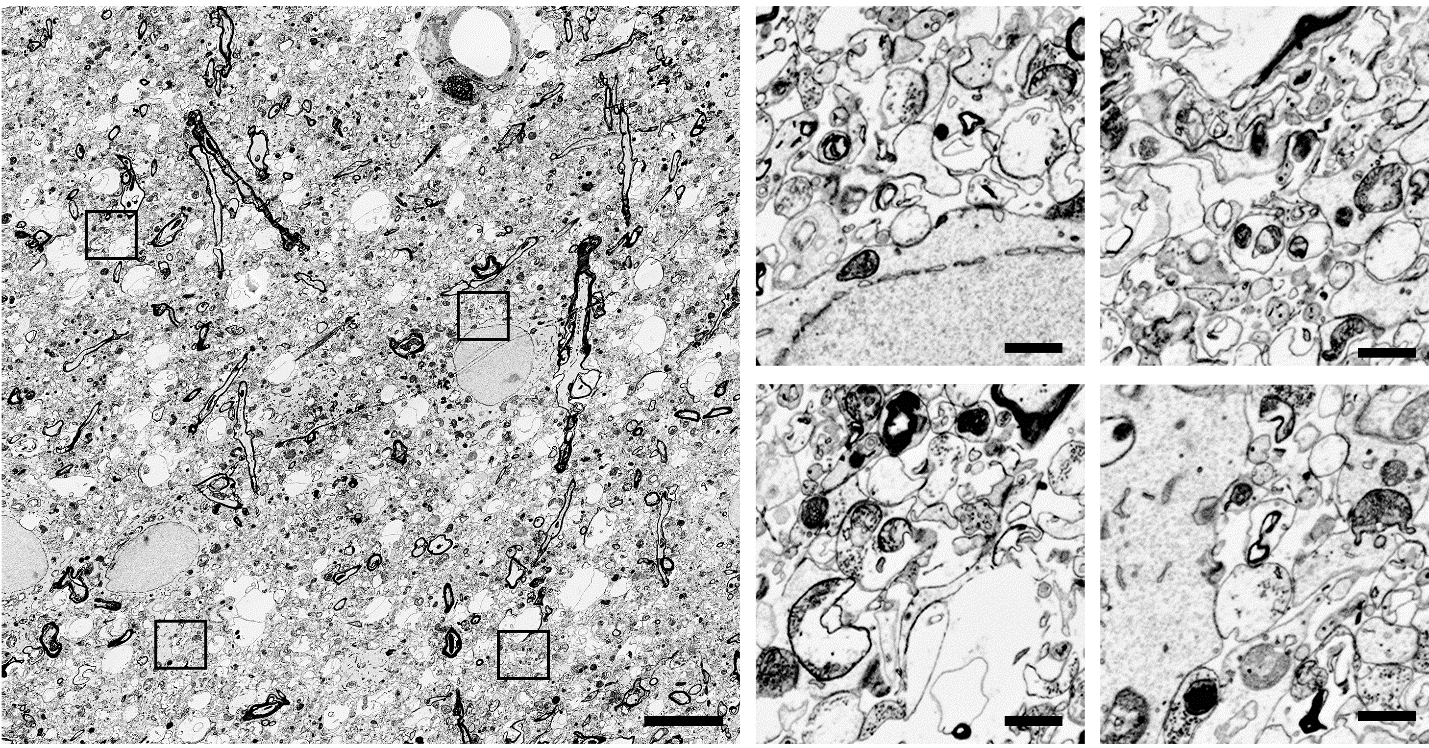

**Supplementary Figure 1.** Electron microscopy imaging of candidate membrane labels. 100 μM of farnesylated glycine pentalysine peptide was applied to 200μm thick tissue slices from a mouse brain perfused with 4% PFA and 0.1% glutaraldehyde in 4⁰C, labelled with lipids for 2 days at 4⁰C and labelled with 0.8nm undecagold gold nanoparticles. In this case, and to achieve the smallest possible probe, we used the azide versions of the probe and conjugated it to dibenzocyclooctyne modified gold nanoparticles. The tissue was post-fixed in 2% glutaraldehyde, embedded in resin, counter-labeled with osmium tetraoxide and imaged. Osmium tetraoxide labeling appears as a grey outline of membranes whereas the lipid labels appear as darker black lines. The farnesylated version of the label only shows partial coverage. Scale bars: 10 μm for the low-resolution images, 1 μm for the high- resolution inserts.

**Supplementary Figure 2.**

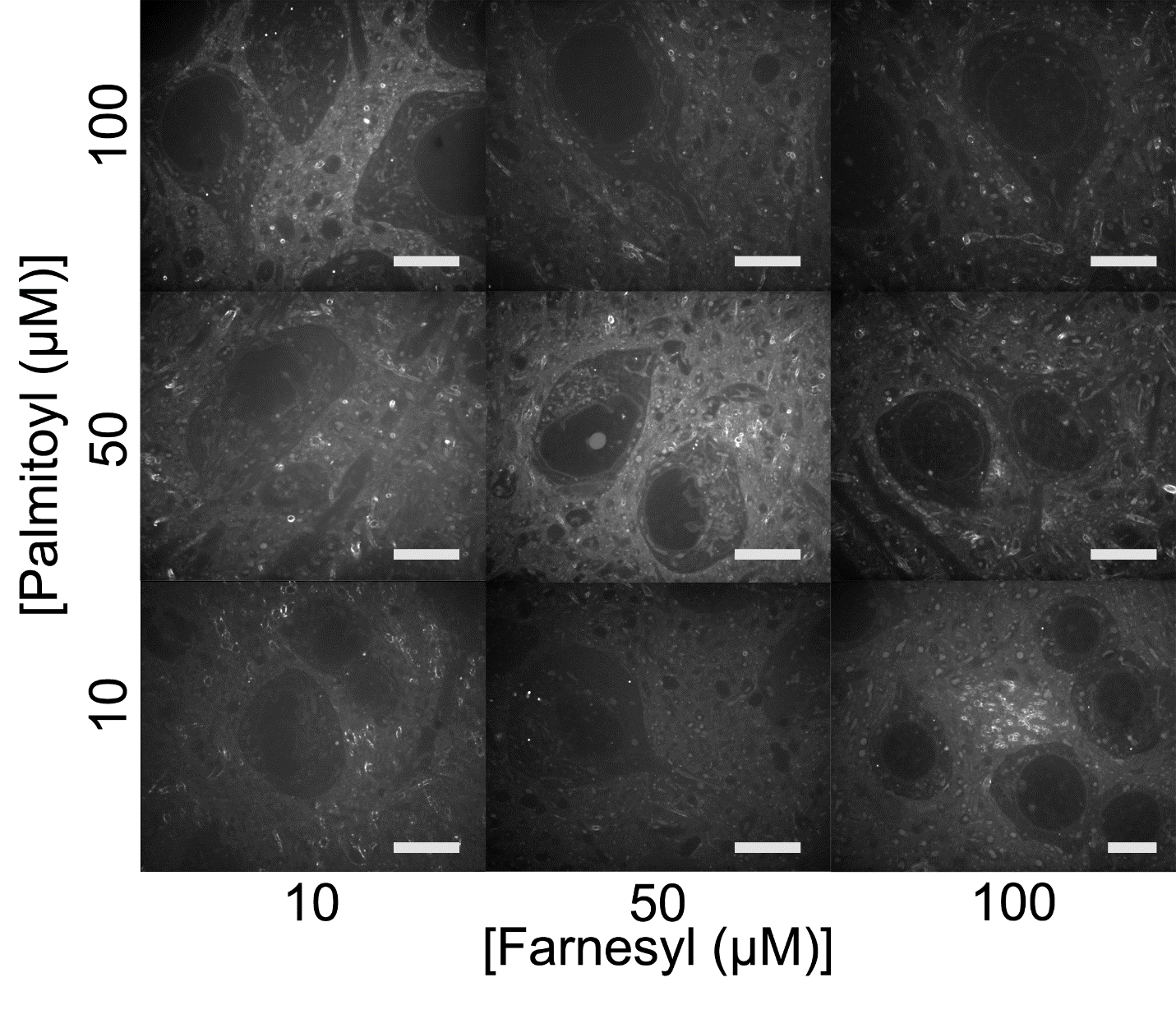

**Supplementary Figure 2.** Combinatorial screening of saturated (palmitoyl) and unsaturated (farnesyl) lipid peptides for membrane labeling. 100μm thick brain tissue, fixed with 4% PFA and 0.1% glutaraldehyde, was incubated at varying concentrations of membrane label containing the two different lipid tails. The tissue was processed with AcX and a bis-acrylamide gel was formed. After tissue homogenization with proteinase K the membrane labels were stained with fluorescent streptavidin, expanded, and imaged. We examined whether combining saturated and unsaturated lipids at various concentrations would increase the membrane labeling yield, but did not observe a major difference. Especially at high concentrations, when combining the two probes, we observed an apparent weaker signal which may be due to either aggregation of the probes due to hydrophobicity or the formation of nanoparticles by them which hinders their diffusion. Scale bars represented in pre-expansion units: 10μm.

**Supplementary Figure 3.**

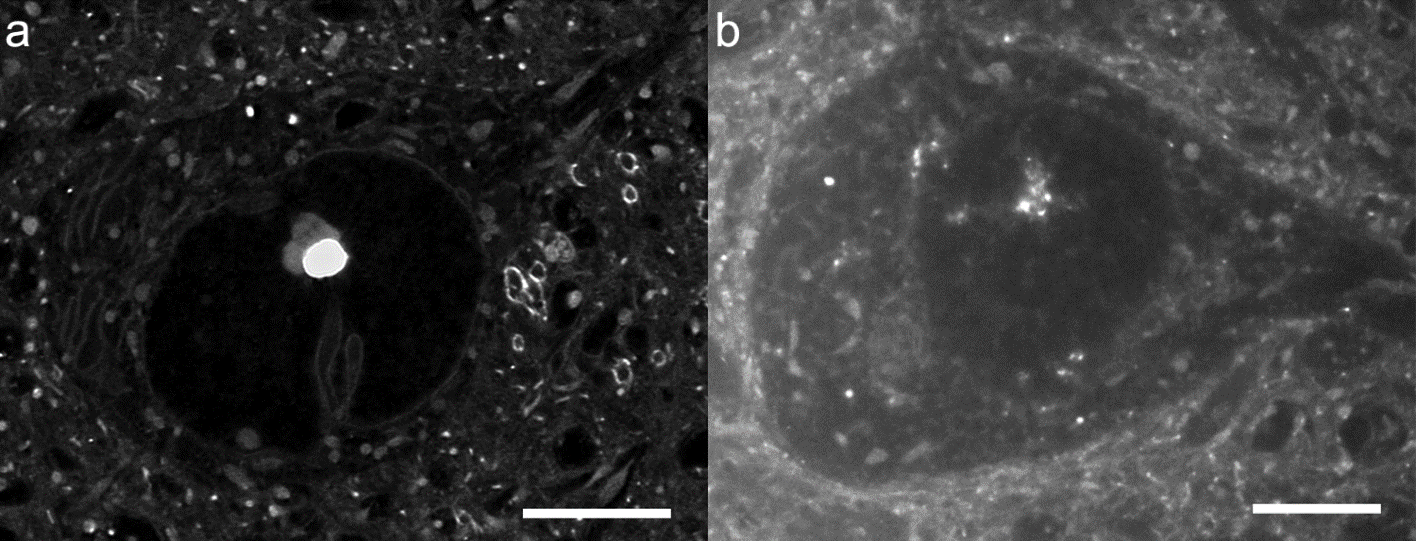

**Supplementary Figure 3.** Effect of the glycine linker attached to the palmitoyl group on the efficacy of membrane labeling in fixed brain tissue. We tested two versions of the palmitoylated penta-lysine biotin membrane probe: a) one containing a glycine linker attached to the palmitoyl group enabling flexibility of the lipid relative to the peptide carrier and b) one that does not contain a glycine but in which the lipid is directly attached to the lysine backbone. In the case of the glycine linker, the level of detail we achieve in labeling membranes is superior to that achieved without the glycine linker. Scale bars represented in pre-expansion units: 5μm.

**Supplementary Figure 4.**

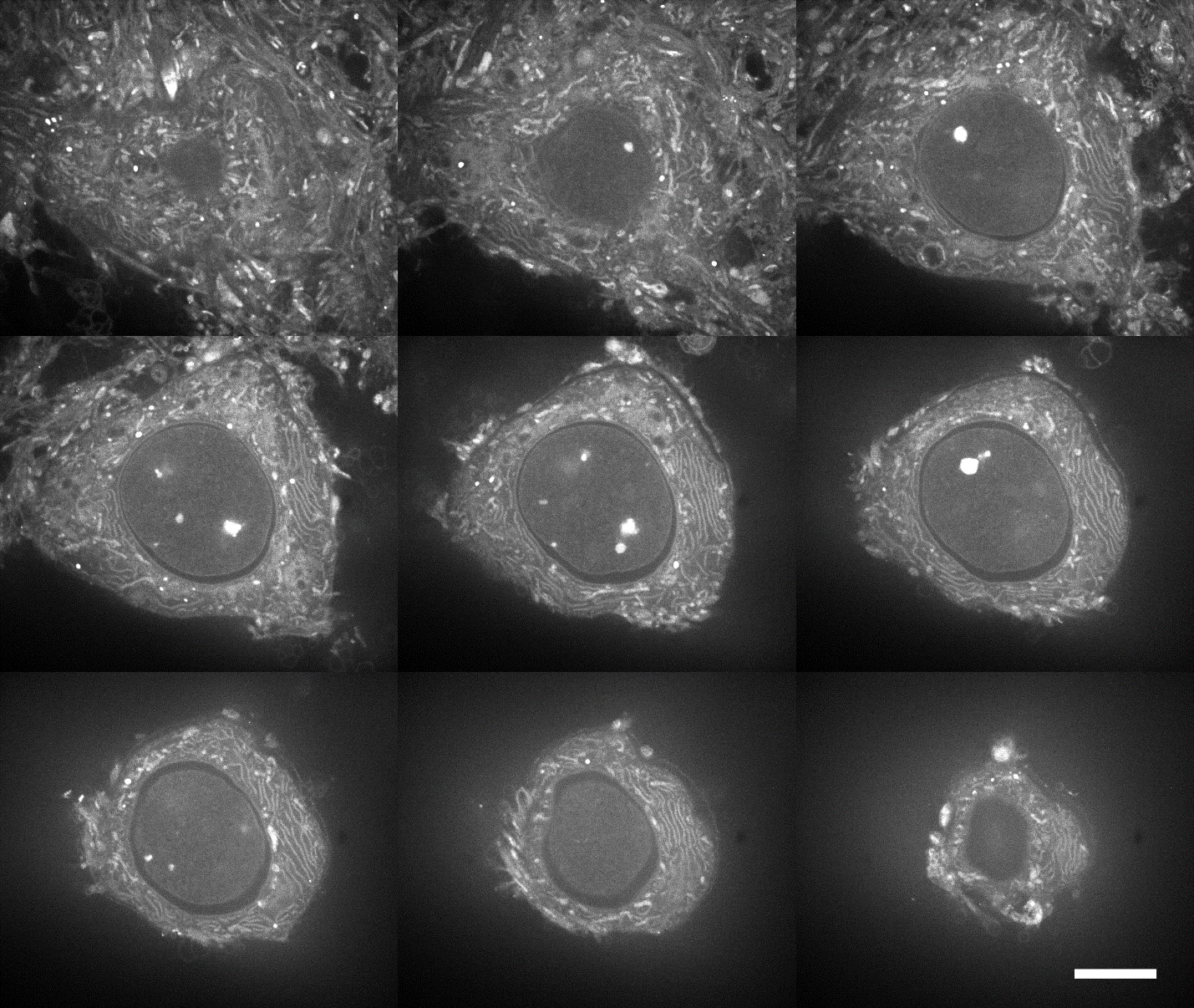

**Supplementary Figure 4.** Membrane labeling of a fixed HeLa cell cultured in vitro. HeLa cells were grown on coverslips and fixed with ice cold 4% PFA and 0.1% glutaraldehyde for a half hour in 4⁰C. The cells were washed overnight with PBS at 4⁰C and 1μM of pGk5b was applied in PBS at 4⁰C for 6 hours. After applying AcX at 0.1mg/ml in PBS for 8 hours at 4⁰C, a bis-acrylamide ExM gel was formed (as in the tissue processing form of mExM) and the samples were processed with proteinase K at 37⁰C for 8 hours. The samples were labeled with streptavidin at 0.1mg/ml concentration and then with fluorescent biotin at 0.2mg/ml. After expansion in water the samples were imaged with a spinning disk confocal microscope. Nine different z-sections of a single stack captured 0.5μm apart are displayed. We can achieve labeling of cellular organelles and membranes and image them with nanoscale resolution. Scale bar represented in pre-expansion units: 5μm.

**Supplementary Figure 5.**
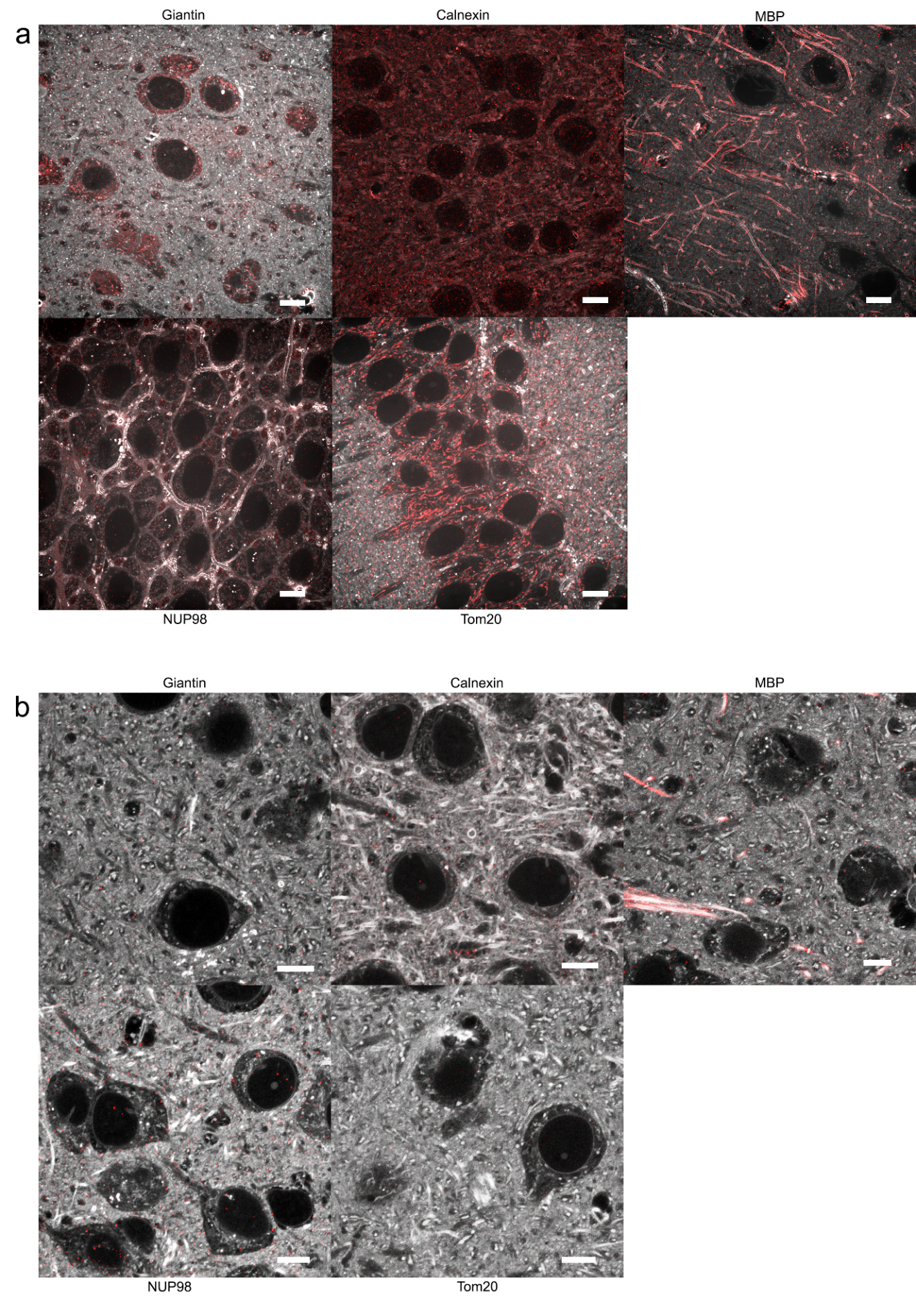

**Supplementary Figure 5.** Attempst to establish an antibody labeling procedure by synthesizing chemical meshes in an already-membrane labeled tissue. Considering that membrane and antibody labeling are not directly compatible, as during antibody labeling detergents dissolve lipid membranes, we tried to create a protocol for antibody labeling where after the membrane label application we create a chemical mesh holding those labels in place while performing common immunohistochemistry procedures. First (a) we applied the membrane labels to a 4% PFA and 0.1% glutaraldehyde fixed tissue and post-fixed with 0.1% PFA. After antibody labeling following established immunohistochemistry protocols including permabilization with detergents, we gelled the tissue, homogenized it with proteinase K and expanded. The lipids are colorcoded with black and white, and the antibody signal with red. We observed that not all of the antibodies labeled the epitopes correctly, for example calnexin labeling becomes non specific (white arrows; the signal is non-specific) and NUP98 labeling is lost (white arrows; no signal on the nuclear membrane), and the tissue expanded only ~3-fold. Alternatively (b), after applying the membrane labels we created a non-expanding hydrogel, labeled with antibodies, created a second expanding hydrogel on top and after homogenization with proteinase K expanded the tissue. In that case we observed loss of labeling with antibodies and the tissue also expanded 3-fold. The loss of labeling may be due to the decrease of diffusivity of the antibodies imposed by the added non-expanding hydrogel. Scale bars represented in pre-expansion units: 10μm.

**Supplementary Figure 6.**

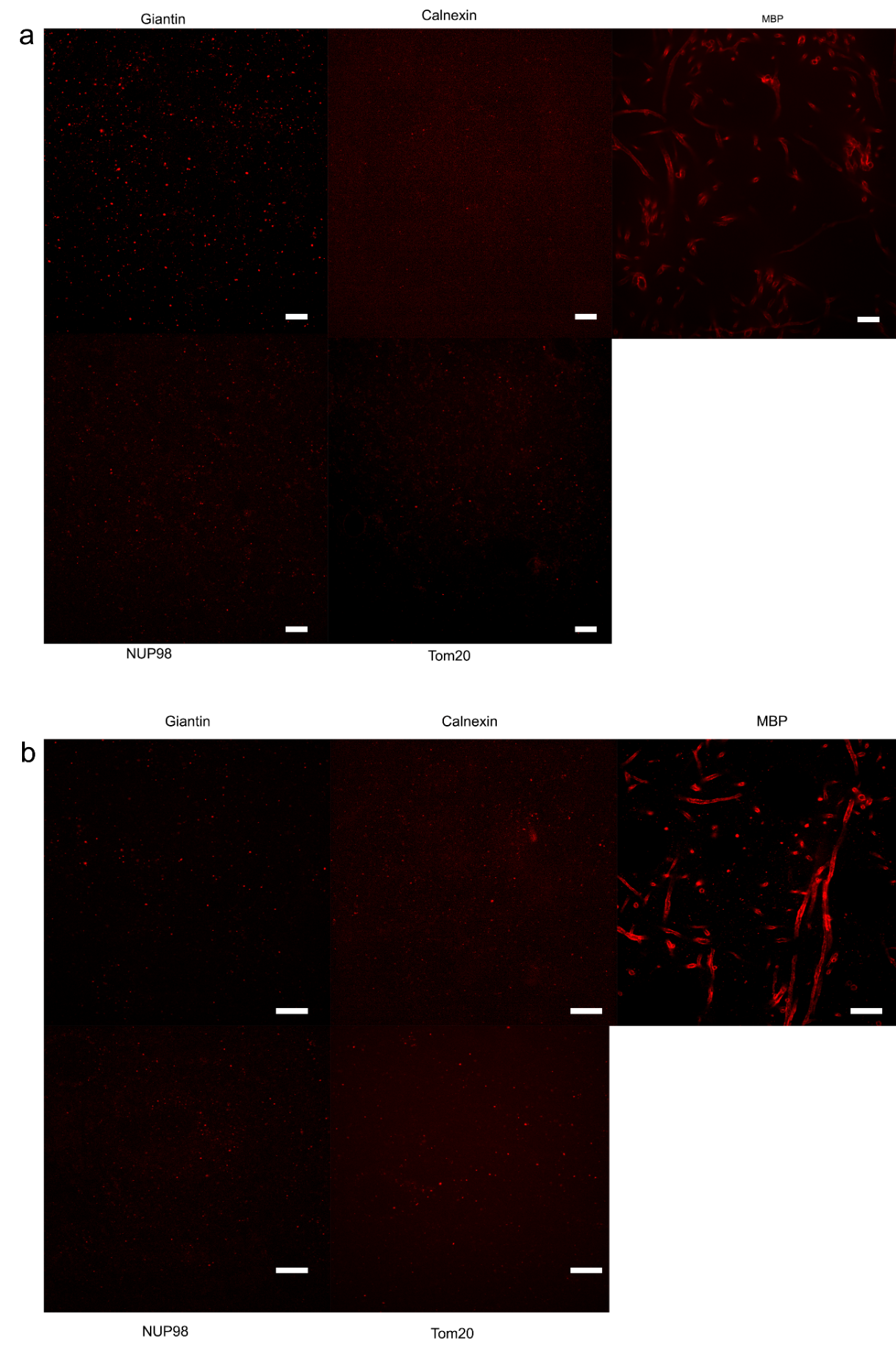

**Supplementary Figure 6.** Attempts to establish an antibody labeling procedure by processing the membrane labeled and gelled tissue with common antigen retrieval buffers. Fixed tissue was labeled for membranes with pGk5b and gelled. The gel-embedded tissue was thermally processed in a sodium citrate buffer (10mM Sodium Citrate, 0.05% Tween 20, pH 6.0) (a) or tris buffer (10mM Tris Base, 0.05% Tween 20, pH 10) (b), autoclaving at 121⁰C for 1 hour. The gels were later labeled with antibodies and imaged. In both cases, we observed loss of antibody signal for most of the tested antibodies. Scale bars represented in pre-expansion units: 10μm.

**Supplementary Figure 7.**

Microscope Slide

Coverslip #1.5

Polymerization Solution

Parafilm

Strip

Tissue

**Supplementary Figure 7.** The tissue polymerization chamber. To achieve polymerization, the tissue is set submerged in polymerization solution in between two pieces of glass: a microscope slide that serves as the base of the polymerization chamber, and a coverslip (thickness #1.5). To prevent compression of the tissue, the coverslip is set onto two strips of parafilm, which also prevent movement of the coverslip or leaking of the polymerization solution.

**Supplementary Table 1.**

**Monomer solution aka StockX (9.4ml, aliquoted to 10 tubes of 940μl and stored at -20⁰C):**

| **Reagent** | **Stock concentration*** | **Amount (ml)** |
| --- | --- | --- |
| Sodium Acrylate (Sigma, cat. no. 408220) | 38 | 2.25 |
| Acrylamide (Sigma, cat. no. A8887) | 50 | 0.5 |
| N,N′-Methylenebisacrylamide (Sigma, cat. no. M7279) | 2 | 0.75 |
| Sodium chloride (Thermo Fisher, cat. no. BP358-212) | 29.2 | 4 |
| PBS (Thermo Fisher, cat. no. 70011044) | 10x | 1 |
| Water (Thermo Fisher, cat. no. 10977015) |  | 0.9 |
| **Total** |  | **9.4** |

*All concentrations are in g/100 ml except PBS. All stock solutions are formulated in water (Thermo Fisher, cat. no. 10977015).

**Polymerization solution (1ml, prepared at 4⁰C, gelled at 37⁰C):**

| **Reagent** | **Stock concentration*** | **Amount (μl)** |
| --- | --- | --- |
| Monomer Solution (see above) | 1x | 940 |
| 4-hydroxy-TEMPO (Sigma, cat. no. 176141) | 0.5 | 20 |
| TEMED (Sigma, cat. no. T7024) | 10 | 20 |
| APS (Thermo Fisher, cat. no. 17874) | 10 | 20 |
| **Total** |  | **1000** |

*All concentrations are in g/100 ml except Monomer Solution. All stock solutions are formulated in water (Thermo Fisher, cat. no. 10977015).

**Digestion buffer* (100ml, prepared and stored at RT, applied at 37⁰C):**

| **Reagent** | **Stock concentration** | **Amount** |
| --- | --- | --- |
| Tris pH 8.0 (Thermo Fisher, cat. no. AM9856) | 1M | 5ml |
| EDTA (Thermo Fisher, cat. no. 15575020) | 0.5M | 0.2ml |
| Triton X-100 (Sigma, cat. no. X100) | 10% | 5ml |
| NaCl (Sigma, cat. no. S5886) | >99% solid | 5.85g |
| Water (Thermo Fisher, cat. no. 10977015) |  | 84ml |
| **Total** |  | **100ml** |

*To formulate the Digestion solution, dilute Proteinase-K (NEB, cat. no. P8107S) at 1:100 dilution in Digestion buffer. All stock solutions are formulated in water (Thermo Fisher, cat. no. 10977015).

**Fixation Reversal buffer (10ml, prepared at RT and used immediately):**

| **Reagent** | **Stock concentration*** | **Amount** |
| --- | --- | --- |
| PEG20000 (Sigma, cat. no. 95172-250G-F) | 5% | 1ml |
| DTT (Thermo Fisher, cat. no. R0862) | >97% solid | 154.3mg |
| SDS (Thermo Fisher, cat. no. AM9820) | 20% | 2ml |
| Tris pH8 (Thermo Fisher, cat. no. AM9856) | 1M | 1ml |
| Water (Thermo Fisher, cat. no. 10977015) |  | 5.9ml |
| **Total** |  | **10ml** |

*****All stock solutions are formulated in water (Thermo Fisher, cat. no. 10977015).

**Supplementary Table 2.**

| Antigen | Species | Company | Catalog no | Boiling (0.5h@100⁰C, 2h@80⁰C) | Autoclaving  (1h@121⁰C) |
| --- | --- | --- | --- | --- | --- |
| Calnexin | Rabbit | Abcam | ab22595 | ✓ | ✓ |
| Tom20 | Rabbit | CST* | 42406S | ✓ | ✓ |
| Tom20 | Mouse | SCBT** | sc-17764 | ✓ | 🗶 |
| NUP98 | Rabbit | CST | 2597S | ✓ | ✓ |
| MBP | Rabbit | CST | 78896S | ✓ | ✓ |
| MBP | Chicken | AVES | AB_2313550 | ✓ | ✓ |
| Giantin | Rabbit | Biolegend | 924302 | ✓ | 🗶 |
| Calreticulin | Rabbit | CST | 12238S | ✓ | 🗶 |

* Cell Signaling Technology

**Santa Cruz Biotechnology

✓: Working

🗶: Not working

**Supplementary Materials and Methods**

**Electron microscopy of lipid labeled tissue**

A terminally isoflurane anesthetized mouse was transcardially perfused with 1x phosphate buffered saline (PBS) at 4⁰C until the blood cleared, followed by 4% PFA and 0.1% glutaraldehyde at 4⁰C. The tissue was post-fixed in the same solution (4% PFA and 0.1% glutaraldehyde in 4⁰C) overnight. The tissue was sectioned into 200μm thick slices using a vibratome (Leica VT1000 S) and washed in PBS, in 4⁰C for a week. For each tissue section, 100 μM of palmitoylated glycine pentalysine peptide or 100 μM of farnesylated glycine pentalysine peptide was applied for 2 days at 4⁰C and washed overnight in PBS at 4⁰C. In this case, and to achieve the smallest possible probe, we used the azide versions of the probes (i.e., the C-terminus of the peptide was modified with an azide). The membrane probes were post-labelled with 0.8nm undecagold gold nanoparticles modified with a dibenzocyclooctyne. Briefly 50nmole of amine modified 0.8nm undecagold gold nanoparticles (Nanoprobes) diluted in PBS were incubated with 500nmole of DBCO-NHS ester (Click Chemistry Tools) dissolved in DMSO overnight at room temperature. After the conjugation reaction the solution was further diluted to achieve <0.1% DMSO v/v in PBS and the reaction product was purified with dialysis in a membrane with 500 Da MWCO (SpectraPor/Spectrum). The final product solution was concentrated in the dialysis membrane after submerging the membrane in a hygroscopic solution of 40%(w/v) PEG20,000 (Sigma Aldrich) in water to final volume 1ml. The DBCO modified gold nanoparticles in PBS solution were applied to the membrane labeled tissue for 2 days at 4⁰C. The tissue was post-fixed in 2% glutaraldehyde, embedded in resin, counter-labeled with osmium tetraoxide and imaged. The tissue processing and imaging, post-membrane labeling, was performed at the electron microscopy facility at the Center of Brain Science at Harvard. Briefly, the tissue was counter-stained with reduced osmium tetroxide-thiocarbohydrazide-osmium and infiltrated with Epon resin. After curing, the tissue was sectioned to 30nm thick slices using a commercial ultramicrotome (ATUM) and imaged with a Sigma scanning electron microscope (Carl Zeiss), equipped with the ATLAS software (Fibics).

**Antibody labeling in a “non-expanding” gel**

A terminally isoflurane anesthetized mouse was transcardially perfused with 1x phosphate buffered saline (PBS) at 4⁰C until the blood cleared, followed by 4% PFA and 0.1% glutaraldehyde at 4⁰C. The tissue was post-fixed in the same solution (4% PFA and 0.1% glutaraldehyde in 4⁰C) overnight. The tissue was sectioned into 100μm thick slices in a vibratome (Leica VT1000 S) and washed in PBS, in 4⁰C for a week. Tissue slices were incubated with the pGk5b lipid label at 4⁰C overnight. Subsequently, 6-((acryloyl)amino)hexanoic Acid, Succinimidyl Ester (AcX) stock solution (10 mg/mL in dimethylsulfoxide (DMSO)) was diluted in 1x PBS that was kept at 4⁰C at a 1:100 dilution, and the tissue slices were incubated in the diluted AcX solution. A non-expanding cleavable hydrogel was cast in the tissue. To prepare 9.4ml of this gel monomer solution, we combined 2ml of 50%(w/v) acrylamide (Sigma Aldrich) in water, 1.5ml of 5%(w/v) N,N'-diallyl-tartardiamide (Bio-Rad) in water, 1ml of 10x PBS and 4.9 ml of water. To initiate the polymerization reaction, to the 9.4ml of monomer solution we added 200μl of 0.5%(w/v) 4-hydroxy-TEMPO in water, 200μl of 10%(w/v) TEMED in water and 200μl of 10%(w/v) APS also in water. The monomer solution was kept on ice and incubation of the tissue slices in the monomer solution was performed at 4⁰C for 30min. Following the established ExM protocol the gel was formed in situ at 37⁰C. The sample was permeabilized in 1x PBS, 0.3% Triton-X buffer for 2 hours at room temperature and then blocked in blocking buffer (1x PBS, 5% Normal Donkey Serum, 0.3% Triton-X) for 4-6 hours also at room temperature. Antibodies were diluted according the above Materials and Methods section in antibody dilution buffer (1x PBS, 1% Normal Donkey Serum, 0.3% Triton-X) and applied to the samples for 10-12 hours at room temperature. After washing for 4 hours in 1x PBS, 0.3% Triton-X and changing the washing buffer every 30 minutes, secondary antibodies were added in the same antibody dilution buffer for 10-12 hours at room temperature and at the same concentration as already stated in the Materials and Methods above. The membrane labels were counter-stained with fluorescent streptavidin similarly to the mExM protocol and washed for 2 hours at room temperature with PBS. Subsequently AcX (final concentration 0.1mg/ml in PBS) was applied to the samples at 4⁰C overnight. A new non-cleavable expanding gel was formulated on top of the cleavable non-expanding gel. For that, Stock X solution was formulated similarly to the Materials and Methods above and 4HT/TEMED/APS was added. The samples were incubated with that solution for 45 minutes at 4⁰C and gelled according to the procedure mentioned in the Materials and Methods above. The double gelled samples were digested with proteinase K as above, overnight at 37⁰C. The first non-expanding gel was cleaved by incubating the samples in 20 mM sodium periodate (Sigma Aldrich, cat# 71859), pH 5.5 in PBS, for 30 minutes at 37⁰C. The gel was subsequently washed in PBS, overnight at 4⁰C, and the signal from the membrane labels was amplified by labelling the gel once more with fluorescently labelled biotin as mentioned in the Materials and Methods above. After expanding the final gel in water, imaging was performed.

**Antibody labeling with double fixation**

100 μm thick tissue slices were first incubated in the lipid labels (e.g., pGk5b) at 4⁰C overnight to let the labels diffuse and intercalate thoroughly throughout the tissue slices. The lipid labels are stored at -20⁰C in 10mM stock concentration (in 50/50 water/DMSO mix), and they are diluted in 1x PBS that is kept at 4⁰C at 1:100 dilution for incubating the 100 μm thick tissue slices. After brief washing in PBS for 2 hours at 4⁰C, the tissue was fixed again in 0.1% PFA solution in PBS at 4⁰C for 4 hours and washed in PBS overnight at 4⁰C. The sample was permeabilized in 1x PBS, 0.3% Triton-X buffer for 2 hours at room temperature and then blocked in blocking buffer (1x PBS, 5% Normal Donkey Serum, 0.3% Triton-X) for 4-6 hours also at room temperature. Antibodies were diluted according the above Materials and Methods section in antibody dilution buffer (1x PBS, 1% Normal Donkey Serum, 0.3% Triton-X) and applied to the samples for 10-12 hours at room temperature. After washing for 4 hours in 1x PBS, 0.3% Triton-X and changing the washing buffer every 30 minutes, secondary antibodies were added in the same antibody dilution buffer for 10-12 hours at room temperature and at the same concentration as already stated in the Materials and Methods above. The membrane labels were counter-stained with fluorescent streptavidin similarly to the mExM protocol and washed for 2 hours at room temperature with PBS. The gelled samples were digested with proteinase K as above, overnight at 37⁰C. They were washed in PBS for 2 hours at room temperature. The signal from the membrane labels was amplified by labelling the gel once more with fluorescently labelled biotin as mentioned in the Materials and Methods above. After expanding the final gel in water, imaging was performed.

**Antibody labeling with saponin**

100 μm thick tissue slices were first incubated in the lipid labels (e.g., pGk5b) at 4⁰C overnight to let the labels diffuse and intercalate thoroughly throughout the tissue slices. The lipid labels are stored at -20⁰C in 10mM concentration (in 50/50 water/DMSO mix), and they are diluted in 1x PBS that is kept at 4⁰C at 1:100 dilution for incubating the 100 μm thick tissue slices. After brief washing in PBS for 2 hours at 4⁰C, the sample was first permeabilized in 1x PBS, 0.1% saponin for 2 hours at room temperature and then blocked in blocking buffer (1x PBS, 5% Normal Donkey Serum, 0.1% saponin) for 4-6 hours also at room temperature. Antibodies were diluted according the above Materials and Methods section in antibody dilution buffer (1x PBS, 1% Normal Donkey Serum, 0.1% saponin) and applied to the samples for 10-12 hours at room temperature. After washing for 4 hours in 1x PBS, 0.1% saponin and changing the washing buffer every 30 minutes, secondary antibodies were added in the same antibody dilution buffer for 10-12 hours at room temperature and at the same concentration as already stated in the Materials and Methods above. The membrane labels were counter-stained with fluorescent streptavidin similarly to the mExM protocol and washed for 2 hours at room temperature with PBS. The gelled samples were digested with proteinase K as above, overnight at 37⁰C. They were washed in PBS for 2 hours at room temperature. The signal from the membrane labels was amplified by labelling the gel once more with fluorescently labelled biotin as mentioned in the Materials and Methods above. After expanding the final gel in water, imaging was performed.
